# Asymmetric dimethylation of Ribosomal S6 Kinase 2 regulates its cellular localisation and pro-survival function

**DOI:** 10.1101/2022.10.18.512681

**Authors:** Mahmoud I. Khalil, Heba M. Ismail, Ganna Panasyuk, Ivan Gout, Olivier E. Pardo

## Abstract

Ribosomal S6 Kinases (S6Ks) are critical regulators of cell growth, homeostasis, and survival, with dysregulation of these kinases being associated with various malignancies. While S6K1 has been extensively studied, S6K2 has been neglected despite its reported involvement in cancer progression. Protein arginine methylation is a widespread post-translational modification regulating a plethora of biological responses in mammalian cells. Here we report that p54-S6K2 is asymmetrically dimethylated at Arg-475 and Arg-477, two conserved residues within the AT-hook motif of the S6K2 family and some AT-hook-containing proteins. We demonstrate that PRMT1, PRMT3, and PRMT6 bind to and methylate S6K2 *in vitro* and *in vivo*. This methylation localises S6K2 to the nucleus where it rescues cells from starvation-induced cell death. Taken together, our findings highlight a novel mechanism regulating the biological function of p54-S6K2 that may be relevant to cancer where Arg-methylation is often found elevated.

## INTRODUCTION

Ribosomal protein S6 kinases (S6Ks) are serine/threonine kinases which are downstream effectors of the extracellularly regulated kinases 1/2 (ERK1/2), the phosphoinositide 3-kinase (PI3K) and the mammalian target of rapamycin (mTOR) [1]. Despite their sequence similarity, S6K1 and 2 are involved in distinct cellular processes and many studies have shown different regulatory mechanisms for S6K2 [2–5]. S6Ks are encoded by two genes; *RPS6KB1* and *RPS6KB2* encoding S6K1 (p85, p70-, and oncogenic p31-S6K1 isoforms [1]) and S6K2 (p56- and p54-S6K2 isoforms), respectively. The longer isoforms of S6K1 and S6K2 (p85-S6K1 and p56-S6K2) contain 23 and 13 amino acid N-terminal extensions, respectively, that are absent in the shorter isoforms and contain a nuclear localisation signal (NLS). S6K2 has an additional NLS and a proline-rich domain in its C-terminal domain [6]. S6K2 interacts with multiple proteins in the nucleus [3] and its nuclear localisation provides it with a distinct role from S6K1 in cell division and transcriptional regulation [3]. S6K2 has increased nuclear localisation in human breast cancer [7] and has been linked to pro-survival and chemoresistance signalling in various cancers through the phosphorylation of the nuclear RNA-binding protein hnRNPA1 [2].

Protein arginine methylation is a post-translational modification (PTM) that is as common, although not as dynamic, as phosphorylation, acetylation, and ubiquitination in mammals where it controls various biological processes [8]. Protein arginine methyltransferases (PRMTs) catalyse the transfer of a methyl group from S-adenosylmethionine (SAM/AdoMet) to arginine within polypeptides. Methylated arginines have lower hydrogen bonding capacity and increased hydrophobicity, thereby modulating interactions with other proteins or nucleic acids [8]. In particular, proteins with certain domains (i.e. Tudor, PHD, and WD40) can selectively bind to methylated arginine. Most proteins targeted by PRMTs possess glycine-arginine rich (GAR), RGG, RG repeats, or RXR sequence motifs [8].

Nine PRMTs, with relatively conserved catalytic core domains exist, classified into three classes (type I, type II, and type III) according to their catalytic activity and mode of methylation [8]. Type I comprises six PRMTs (PRMT1, −2, −3, −4, −6, and −8) and type II two (PRMT5 and −9). PRMT7 is the only type III PRMT and catalyses only monomethylation (MMA) [8]. Although PRMT1 is the predominant member, the lack of any PRMT cannot be compensated by others [9]. PRMTs promote arginine asymmetric and symmetric dimethylation (sDMA and aDMA, respectively), or MMA, although the functional consequences of sDMA versus aDMA are uncertain [8]. While the existence of dedicated arginine demethylases or enzymes that interfere with arginine methylation has been suggested, this has recently been contested [10] so the dynamics of this PTM is currently uncertain.

Here we demonstrate that p54-S6K2 is asymmetrically dimethylated on Arg-475/477. We show that PRMT1, 3, and 6 bind and methylate S6K2, promoting its nuclear localisation and rescue from starvation-induced cell death. Hence, we provide the first evidence that aDMA regulates the biological functions of p54-S6K2.

## RESULTS AND DISCUSSION

S6K2 is mapped to the chromosomal *11q13* locus, commonly amplified in various cancers [11] where it was shown to regulate cell survival [2, 12, 13]. We describe for the first time, the aDMA of S6K2 and the role of this modification in cell survival. Unlike phosphorylation, acetylation, and ubiquitination which have been extensively examined, arginine methylation has been less investigated.

### S6K2 interacts with PRMT1, PRMT3, and PRMT6 *in vivo* and *in vitro*

The extreme C-terminus of human S6K2 shows significant sequence conservation across mammalian species (**Fig. 1A**). Since the p54-S6K2 C-terminus contains two overlapping RXR motifs (RXRXR) previously shown to be targeted by PRMTs [8], we hypothesized that it may be arginine methylated.

**Fig. 1.**
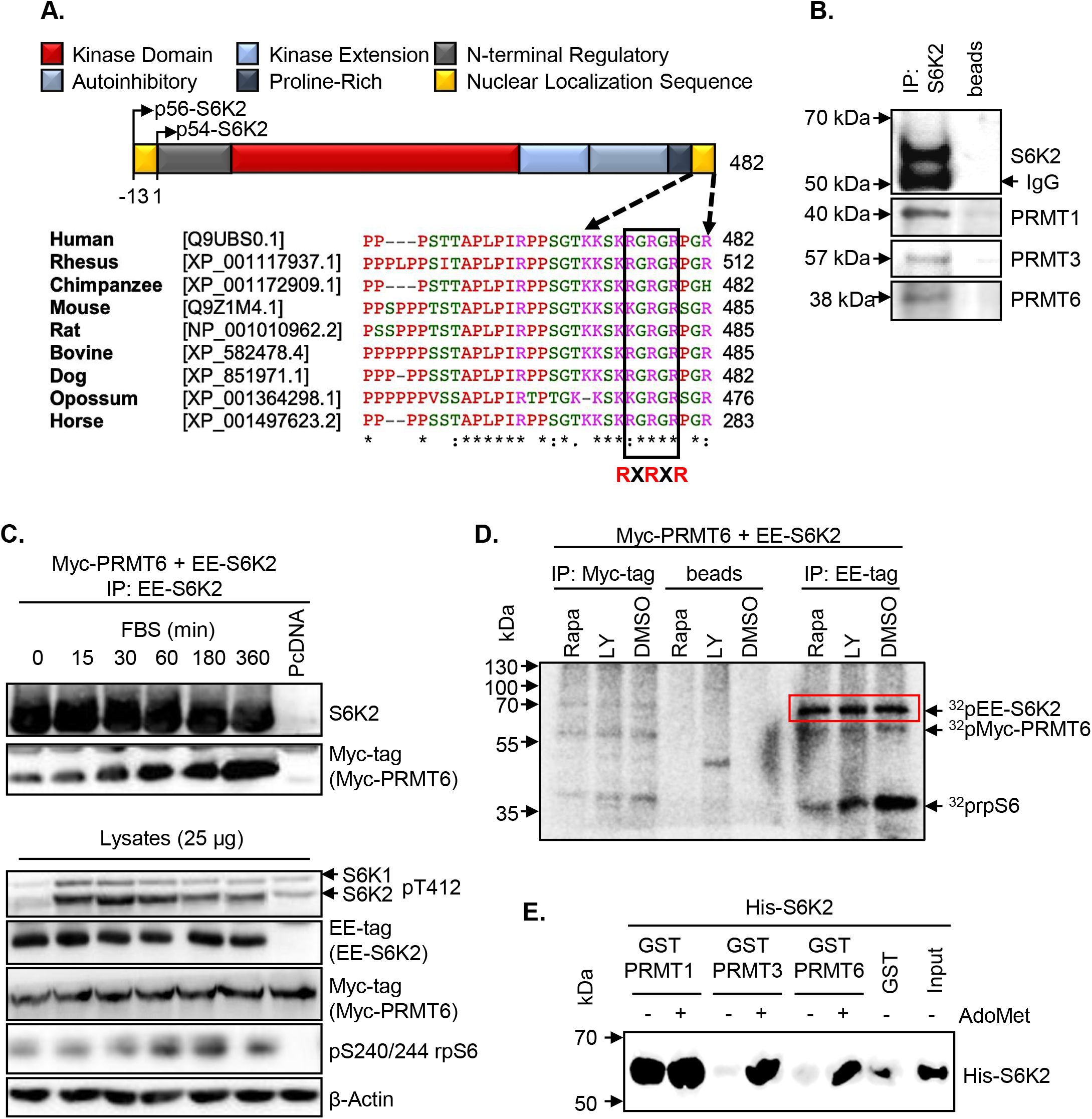
S6K2 interacts with PRMT1, 3, and 6 *in vivo* and *in vitro*. **(A) Protein domain organisation of S6K2 and alignment of the amino acid sequences in its extreme C-terminus**. Conserved Arginine-X-Arginine motif (RXRXR) is boxed, and “X” represents any amino acid. **(B) Endogenous S6K2 interacts with PRMTs (1, 3, and 6) in cells**. S6K2 was immunoprecipitated from HEK293 and interacting proteins were detected by immunoblotting. Protein A-Sepharose beads (beads) alone were used as a control. **(C) The interaction between S6K2 and PRMT6 is induced by serum stimulation.** HEK293 cells expressing Myc-PRMT6 and EE-S6K2 or pcDNA3.1 empty vector (pcDNA) were incubated in serum-free media for 24 h prior to treatment with 10% FBS for the indicated times. Lysates were analysed directly or immunoprecipitated with anti-EE-tag antibody prior to SDS-PAGE/immunoblotting with the indicated primary antibodies. **(D) S6K2 activity is immunoprecipitated with PRMT6**. Lysates from HEK293 cells expressing Myc-PRMT6 and EE-S6K2 were immunoprecipitated with either anti-EE-tag or anti-Myc-tag antibodies following treatment for 30 min with 100 nM Rapamycin (Rapa), 50 μM LY294002 (Ly), or DMSO (vehicle). Immune complexes were subjected to *in vitro* kinase assay using 80S ribosomes as substrate in the presence of [γ-^32^P]ATP prior to SDS-PAGE and autoradiography. Control IP was achieved by incubating lysates with protein A-Sepharose beads (beads). **(E) PRMT1, PRMT3, and PRMT6 interact with S6K2 *in vitro***. GST pull-down assays were performed using His-S6K2 with GST, GST-PRMT1, GST-PRMT3, or GST-PRMT6 in the presence or absence of 200 μM AdoMet, followed by immunoblotting with anti-S6K2 antibody. Input is 0.5 μg His-S6K2. (B-E) All data shown are representative of a minimum of n=3 biological repeats.

Given that interaction of PRMTs with their substrates is required for methylation, we performed co-immunoprecipitation experiments in HEK293 cells which revealed that endogenous S6K2 co-immunoprecipitated with PRMT1, 3, and 6 (**Fig. 1B**). Furthermore, in HEK293 cells overexpressing Myc-tagged PRMT6 and EE-tagged S6K2, this interaction was rapidly (30 min) induced by serum stimulation and steadily increased over 6 hours (**Fig. 1C**). However, kinase activation of S6K2 is not required for this as inhibition of mTOR by Rapamycin or of PI3K by LY294002, both abolishing S6 kinase activity as judged from rpS6 phosphorylation, did not prevent association of S6K2 with PRMT6 (**Fig. 1D**).

To test whether PRMT-S6K2 interaction is direct and whether it is impacted by presence of the methyl donor, His-S6K2 was incubated with GST-PRMT1, 3, or 6 bound to glutathione-Sepharose beads in the presence or absence of the methyl donor, AdoMet. **Figure 1E** shows that AdoMet promotes the interaction of S6K2 with PRMT3 and 6, suggesting that activity or conformational changes of these PRMTs may regulate their binding. In contrast, binding to PRMT1 is constitutive, similar to the previously reported PRMT1/hnRNP interactions [14]. Also, while methylation leaves the overall charge of arginine unchanged, it increases steric hindrance which may disrupt hydrogen bonding [9], modulating protein-protein interactions [15]. S6K2 and PRMTs may regulate each other’s activity through binding, a possibility supported by prior literature suggesting that interactors can either inhibit or activate the methyltransferase activity of PRMTs [16–19].

### S6K2 is methylated *in vitro* and in cells

Given direct interaction between S6K2 and PRMTs, we next tested if S6K2 is a substrate of PRMTs. We performed *in vitro* methylation assays which revealed that full-length S6K2 can be methylated by PRMT1, 3, and 6 (**Fig. 2A**). This was prevented by the SAM-dependent methyltransferase inhibitor, Sinefungin, a competing analogue of AdoMet [8].

**Fig. 2.**
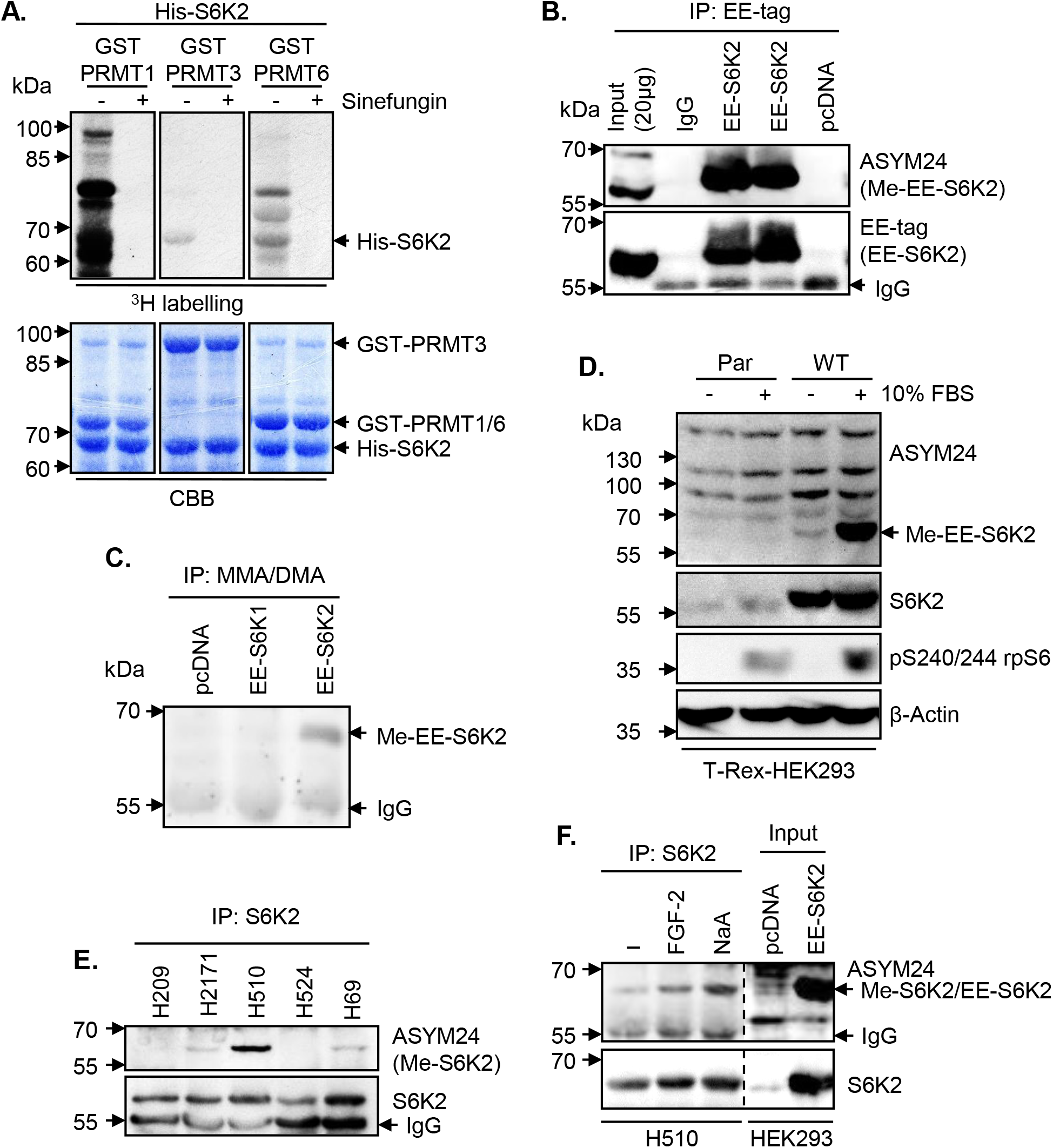
S6K2 is methylated *in vitro* and in cells. **(A) *In vitro* methylation of recombinant S6K2**. *In vitro* methylation assays with His-S6K2 and GST-PRMTs (1, 3, or 6) in the presence of [^3^H]AdoMet, with/without Sinefungin (500 μM). Total amounts of His-S6K2 and GST-PRMTs are shown by Coomassie staining (CBB). **(B) Transiently overexpressed S6K2 is methylated in HEK293 cells**. Lysates from HEK293 cells expressing EE-S6K2 or pcDNA3.1 empty vector (pcDNA) were immunoprecipitated with anti-EE-tag antibody, followed by immunoblotting with the indicated antibodies. IgG was used as immunoprecipitation control. **(C) Methylated EE-S6K2 is immunoprecipitated with MMA/DMA antibody in HEK293 cells**. Lysates from T-Rex-HEK293 cells overexpressing EE-S6K1, EE-S6K2, or pcDNA3.1 were immunoprecipitated with anti-Mono/DiMethyl Arginine (MMA/DMA) antibody, followed by immunoblotting with anti-S6K1 and anti-S6K2 mix of primary antibodies. **(D) S6K2 methylation is induced by serum stimulation**. T-Rex-HEK293 cells overexpressing pcDNA3.1 (Par) or EE-S6K2 (WT) were induced with 1 μg/ml tetracycline for 24 h. Cells were starved in serum-free media (-) for 24 h or stimulated with 10% FBS for 1 h (+) following starvation. Lysates were analysed by SDS-PAGE/immunoblotting with the indicated primary antibodies. **(E) S6K2 is methylated in SCLC cell lines**. Lysates from the indicated SCLC cell lines were subjected to immunoprecipitation with anti-S6K2 antibody, followed by immunoblotting with anti-ASYM24 or anti-S6K2 antibody. **(F) S6K2 methylation in H510 cells is induced by FGF2 and sodium arsenite**. Lysates from untreated (-), 0.1 ng/ml FGF-2-treated (5 min), or 1 mM Sodium arsenite (NaA)-treated (30 min) H510 cells were subjected to immunoprecipitation with anti-S6K2 antibody, followed by immunoblotting with anti-ASYM24 or anti-S6K2 antibody. Input: total lysates from HEK293 cells overexpressing EE-S6K2 or empty vector (pcDNA). (A-F) All data shown are representative of n=3 biological repeats.

All PRMTs tested here are type I enzymes catalysing the formation of asymmetric dimethylarginine (aDMA). To assess whether S6K2 was modified in cells, we used ASYM24, an antibody that specifically recognises aDMA [20]. Ectopically-expressed EE-S6K2 immunoprecipitated from HEK293 cells was detected by ASYM24, confirming that aDMA occurred *in vivo* (**Fig. 2B**). This was confirmed by the detection of EE-S6K2 in MMA/DMA immunoprecipitates from HEK293 cells (**Fig. 2C**). However, this modification was not detected in ectopically-expressed EE-S6K1, showing that this PTM is specific for S6K2, possibly due to amino acid sequence differences in their C-terminus (KSKRGRGRPGR).

As binding of S6K2 to PRMT6 is serum-inducible, we investigated whether S6K2 methylation could be similarly regulated. Immunoprecipitation of transiently expressed EE-S6K2 from HEK293 cells showed that S6K2 methylation was strongly induced following serum stimulation (**Fig. 2D**). The dynamic nature of S6K2 methylation suggests the existence of a negative regulator for this PTM, such as a specific demethylase. Three scenarios for S6K2 demethylation can be proposed to be tested in future work: 1) enzyme-catalysed demethylation [21]; 2) conversion of MMA to citrulline by arginine deiminases preventing subsequent methylation [22]; and 3) replacement of the arginine-methylated pool by unmodified newly-synthesised protein [23].

aDMA-containing proteins have been suggested to play an essential role in oncogenesis, being highly elevated in immortalized cell lines [9]. We previously reported that S6K2, but not S6K1, mediated the survival of small-cell lung cancer (SCLC) cells [2]. As S6K2, but not S6K1, is arginine methylated in our experiments, we wondered if this could be correlated with its pro-survival functions. To test it, we analysed the methylation levels of endogenous S6K2 immunoprecipitated from a panel of SCLC cell lines and found that H2171, H69, and H510 cells showed S6K2 methylation, with the signal in H510 cells being particularly strong (**Fig. 2E**). aDMA was induced in H510 cells treated with FGF-2, which promotes survival in this cell line through S6K2 activation [2], or sodium arsenite, an activator of S6Ks (**Fig. 2F**). Therefore, our results suggest that S6K2 aDMA modification is dynamic. Further testing to identify stimuli inducing or not S6K2 arginine methylation would help understand how widely this PTM participates to signalling specificity.

The interaction of S6K2 with its partners may be regulated by aDMA. Indeed, the two overlapping RXR motifs of S6K2 are preceded by a proline-rich region (PRR), a domain type involved in protein-protein interaction. Interestingly, arginine methylation regulates the binding of PRR to WW and SH3 domain-containing proteins. For instance, hnRNPK arginine methylation near its three PRRs diminishes its association with SRC kinase, reducing its activation and tyrosine phosphorylation of hnRNPK [24]. Similarly, aDMA within the PRR of the RNA-binding protein Sam68 modulates its interaction with SRC [25]. It is possible that SRC-mediated phosphorylation of S6K2 *in vivo* [26] could be affected by arginine methylation. Systematic identification of S6K2 interactors through its PRR would provide deeper understanding of how methylation modulates its function.

### Methylation of S6K2 occurs at Arg-475 and Arg-477

Most PRMTs preferentially methylate substrates within clusters of RGG/RXR box motifs [8] and within Glycine-Arginine-rich (GAR) motifs, present in many DNA- and RNA-binding proteins [27]. While S6K2 contains no GAR motif, it has two overlapping RXR motifs at its extreme C-terminus (RGRGR) encompassing R475, R477, and R479. Interestingly, these three residues are highly conserved in AT-hook domains of some DNA-binding proteins, including the high mobility group (HMG) proteins (**Fig. 3A**). Interestingly, PRMT6 dimethylates HMGA1a on Arg-57/59 in the second and Arg-83/85 in the third AT-hook motif [28], sites equivalent to those identified here for S6K2.

**Fig. 3.**
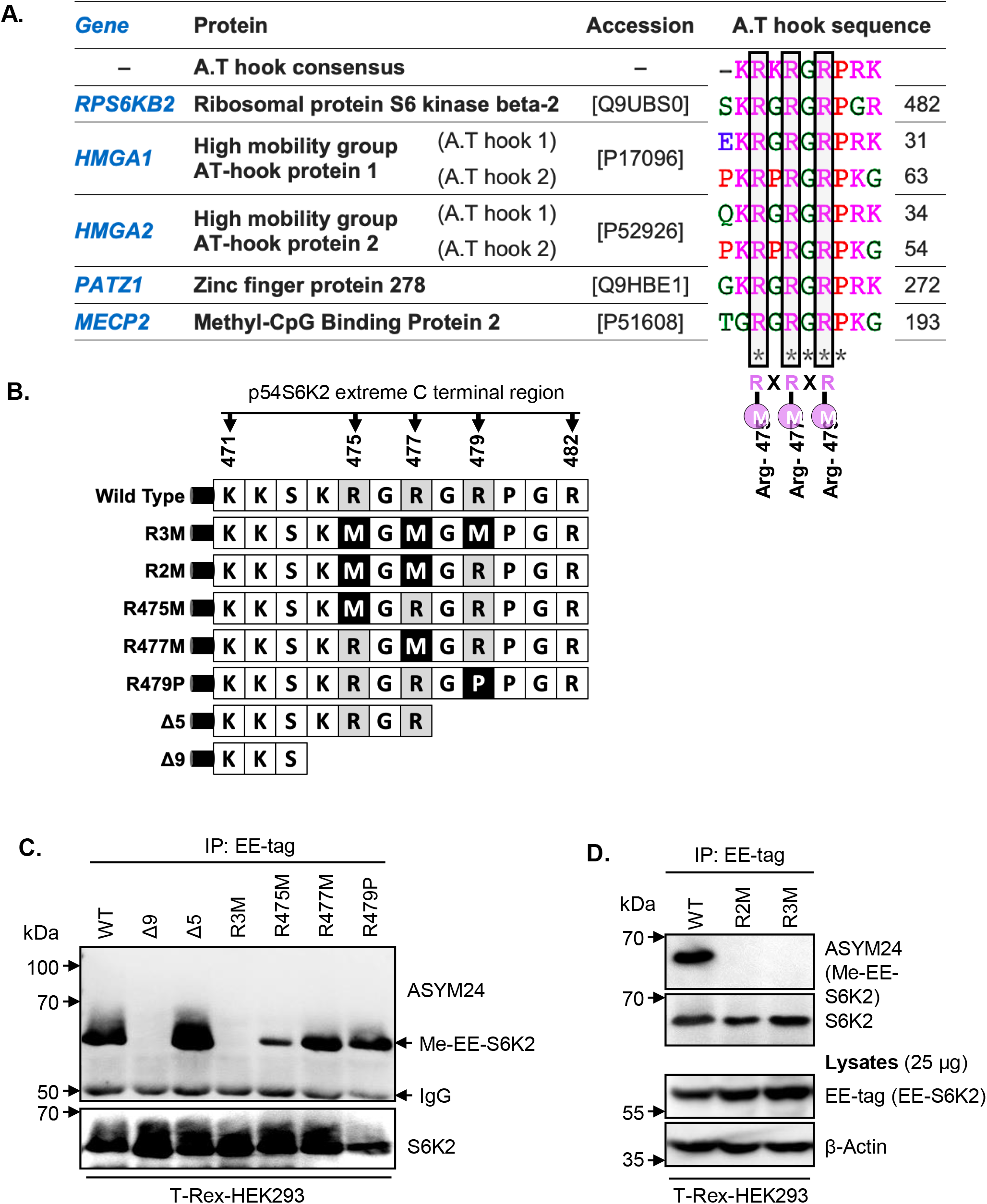
S6K2 is methylated at Arg-475 and Arg-477. **(A) Sequence alignment of the extreme C-terminal region of S6K2 and selected AT-hook-containing proteins**. Putative arginine methylation sites are boxed. RXRXR; consensus sequence for PRMTs substrates. “X” is any amino acid. **(B) Schematic representation of S6K2 mutants**. Single, double, or triple substitution mutants (Arg to Met or Pro) are generated in addition to two truncated mutants (Δ5 and Δ9). **(C-D) S6K2 is dimethylated at Arg-475/477**. Lysates from T-Rex-HEK293 cells stably expressing EE-S6K2 wild type (WT), or different mutants were subjected to immunoprecipitation with anti-EE-tag antibody followed by immunoblotting with the indicated antibodies. **(D)** Total lysates were used as control. All data shown are representative of n=3 biological repeats.

To determine the role of this RXR motif in the dimethylation of S6K2, we mutated Arg-475, Arg-477, and Arg-479 individually and in combination (R475M, R477M, R479P, R475M/R477M (R2M), R475M/R477M/R479M (R3M)) (**Fig 3B**). We also truncated the last 5 or 9 amino acids from the C-terminus (Δ5 and Δ9, respectively). Overexpression in HEK293 cells showed that the Δ5 mutant was methylated similarly to the wild-type protein, while the Δ9 truncation abolished methylation (**Fig. 3C**). This narrowed down the methylation sites to R475 and R477. Notably, all single point mutants were less methylated than the wild-type or Δ5 mutant, suggesting that R479 still modulates this PTM, maybe through stabilising interaction with PRMTs. However, the double (R2M) and triple (R3M) mutants both fully prevented methylation of S6K2, suggesting R475 and R477 as the two major sites modified (**Fig. 3D**).

AT-hook motifs are non-classical DNA-binding domains that interact with short stretches of adenines and thymidines (4-6 bp) centred on the sequence AA(T/A)T in the DNA minor groove. The presence of an AT-hook motif in S6K2 suggests that it may bind DNA, and dimethylation might modulate the affinity or sequence specificity of this binding to impact biological output. Indeed, this is observed for other PRMT targets that harbour nucleotide-binding activity [15]. For instance, the efficiency of PRMT1, 3, and 6 to methylate HMGA1 proteins is markedly reduced upon binding of AT-rich regions to DNA [29]. It is suggested that DNA may physically hinder the accessibility of the AT-hooks to PRMTs due to the rigid conformation of HMGA1 induced by DNA binding. Other studies reported that arginine methylation enhanced the DNA-binding capacity of PRMT substrates. For instance, methylation of DNA polymerase β by PRMT6 increases its DNA binding capacity [30]. Conversely, methylated hnRNPA1 exhibits decreased DNA affinity compared with its unmethylated form [31, 32]. Likewise, methylation of Signal Transducer And Activator Of Transcription 1 (STAT1) decreases its binding to DNA and subsequent function [33, 34]. Hence, the potential for S6K2 to bind DNA and how dimethylation may regulate this should be formally investigated.

### S6K2 methylation modulates its subcellular localisation and pro-survival effects

In view of the above, we wondered if methylation could modify the cellular localisation of S6K2. To test this, we performed subcellular fractionation of HEK293 cells expressing wild-type or arginine-mutant S6K2. This revealed that methylated S6K2 was present in nuclear fraction (**Fig. 4A**), while the R2M mutant appeared mostly cytoplasmic (**Fig. 4B**). This, together with our finding that serum and FGF2 stimulation increases S6K2 methylation (**Fig. 2D and F**), agrees with prior knowledge that S6K2 localises to the cytoplasm of serum-starved cells and translocates to the nucleus upon growth factor stimulation [5]. PRMT-mediated methylation was previously shown to impact the nucleocytoplasmic distribution of its targets, including Sam68, RNA helicase A, and hnRNPA [8, 21, 35]. This can be associated with changes in phosphorylation status, as in the case of STAT6 [36], STAT1 [37] and the Npl3p RNA-binding protein [38]. Hence, methylation may influence or cooperate with other PTMs to regulate protein localisation and activity. In support for this, PP2A, the main phosphatase for S6Ks, is activated by amino acid deprivation [39], a condition that occurs under serum starvation that we found to demethylate S6K2. PP2A is known to bind PRMT1, inhibiting its methyltransferase activity [40]. Hence, activation of PP2A under serum starvation could inactivate S6K2 and simultaneously inhibit the dimethylation of this kinase, excluding it from the nucleus (**Fig. 4D**).

**Fig. 4.**
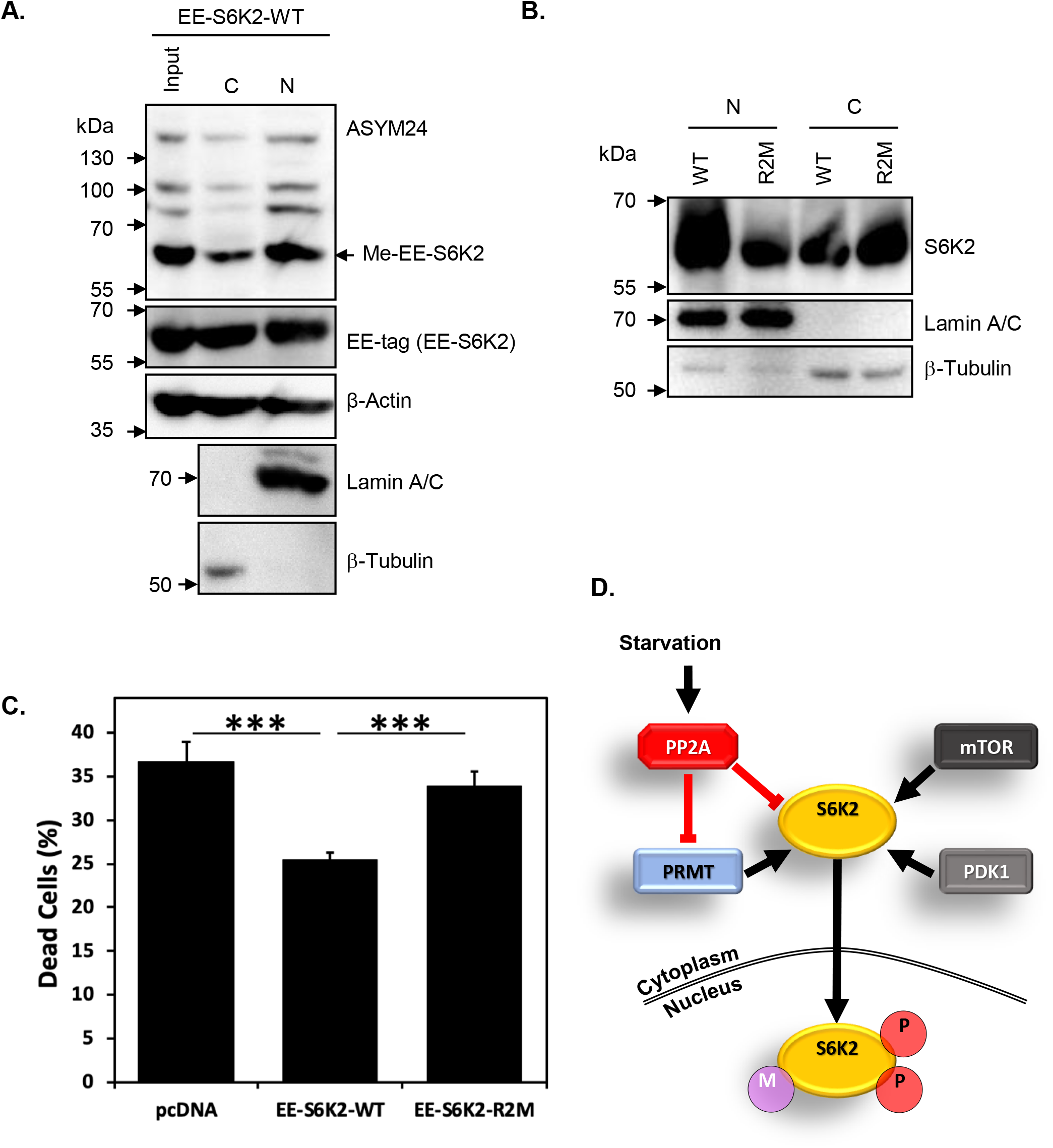
Methylation of S6K2 modulates its subcellular localisation and protects cells from starvation-induced cell death. **(A) Methylated EE-S6K2 is predominantly localised in the nucleus in exponentially growing HEK293 cells**. T-Rex-HEK293 cells stably expressing EE-S6K2 (WT) were fractionated for cytoplasmic (C) and nuclear (N) pools. Whole lysates (input) and fractions were analysed by SDS-PAGE/immunoblotting using the indicated primary antibodies. The purity of fractions was evaluated by blotting with anti-lamin A/C and anti-β-tubulin antibodies. **(B) The R2M mutant is localised in the cytosol of HEK293 cells**. T-Rex-HEK293 cells stably expressing EE-S6K2 (WT) or R2M mutant were fractionated for cytoplasmic (C) and nuclear (N) pools. Fractions were analysed by SDS-PAGE/immunoblotting using the indicated primary antibodies. **(C) Wild type S6K2 (WT), but not R2M mutant, rescues cells from starvation-induced cell death**. HEK293 stably expressing EE-S6K2-WT, EE-S6K2-R2M, or empty vector (pcDNA) were incubated in serum-free media for 24 h. Cells were collected and proportion of dead cells assessed using Trypan blue exclusion (n=12) (\*\*\**p* < 0.001, Student t-test). **(D) Proposed model for regulation of arginine methylation of S6K2**. PRMT, protein arginine methyltransferase; PP2A, protein phosphatase 2A; mTOR, mammalian target of rapamycin: PDK1, 3-Phosphoinositide-dependent kinase 1; P, Phosphorylation; M, arginine methylation. **(A-C)** All data shown are representative of n=3 biological repeats.

We previously demonstrated that S6K2 regulates cell survival through phosphorylation of the nuclear protein hnRNPA1 [2, 13] and showed increased nuclear localisation of this kinase in breast cancer [7] where it drives pro-survival signalling [13]. We therefore hypothesised that arginine methylation of S6K2 may promote cell survival. HEK293 cells expressing either wild-type or R2M mutant S6K2 were serum-starved for 24 hours and cell viability evaluated. As shown in **Fig. 4C**, cells expressing R2M-S6K2 exhibited significantly *(p* < 0.001) increased cell death as compared to cells expressing the wild-type kinase. Taken together, our data suggest that the pro-survival function of S6K2 depends on its dimethylation and positively corelates with its nuclear localization.

Our findings contribute to our understanding of S6K2 activation and biological function. Indeed, PKCε-mediated phosphorylation of receptor-interacting protein 140 (RIP140) stimulated its arginine methylation and subsequent cytoplasmic localisation [41]. Interestingly, PKCε interacts with S6K2 and sequester it in the cytoplasm through phosphorylation of Ser-486, which inhibits the NLS of this kinase [5]. Also, interaction with PKCε is essential for the pro-survival activity of S6K2 in SCLC [2]. Hence, PKCε may regulate S6K2 methylation to modulate its pro-survival activity depending on biological context. In addition, C-terminal phosphorylation by ERK is essential for relieving autoinhibition of S6K2 [3] and we reported that FGF-2 activates S6K2 in SCLC cells through a MEK-dependent pathway [2, 42]. While kinase activity of S6K2 is dispensable for binding to PRMTs (**Fig. 1D**), further work is required to highlight the possible crosstalk between methylation and phosphorylation events and how this may fine-tune S6K2 activity. Indeed, methylation can prevent nearby phosphorylation events, as for the PRMT1-mediated Arg-248/250 dimethylation of FOXO1 which blocks its Ser-253 phosphorylation by AKT, leading to its nuclear accumulation and subsequent induction of apoptosis [43]. Hence, the order in which PTMs occur on S6K2 may define the ultimate biological output obtained.

Taken together, our data reveal an exciting new level of regulation of S6K2 activity that will help clarify how this kinase tailors its biological functions in response to different environmental cues.

## MATERIALS AND METHODS

See Supplementary Information for further details.

## AUTHOR CONTRIBUTIONS

MIK performed the experiments, acquired the data, analysed, and interpreted the data. HMI contributed to the construction of the R3M and truncated mutants of S6K2, and the generation of their stable cell lines. GP contributed to the experiment of the endogenous interaction between S6K2 and PRMTs. MIK, IG, and OEP wrote and edited the manuscript. IG and OEP contributed to the conception and design of this study. All authors have reviewed and approved the final version for submission.

## ACKNOWLEDGEMENTS

We would like to thank A. Bannister and T. Kouzarides (The Gurdon Institute, University of Cambridge, UK) for providing expression plasmids for PRMT1, 3, and 6 and GST fusion proteins. OEP gratefully acknowledge infrastructure support from the Cancer Research UK Imperial Centre, the Imperial Experimental Cancer Medicine Centre and the National Institute for Health Research Imperial Biomedical Research Centre.

## FUNDING

This work was supported by a project grant from the Association for International Cancer Research (AICR 09-0787) and funding from the Cancer Treatment and Research Trust.

## COMPETING INTERESTS

The authors declare no competing interests.

## DATA AVAILABILITY

The data generated during and/or analysed during the current study are available from the corresponding author on reasonable request.

## Notes

### Competing Interest Statement

The authors have declared no competing interest.

