## Supplementary data- Methods for "Asymmetric dimethylation of Ribosomal S6 Kinase 2 regulates its cellular localisation and pro-survival function"

### Supplementary information

#### Supplementary Materials and Methods

##### *General Reagents*

All the general-purpose reagents were purchased from Thermo Fisher Scientific UK Ltd., Sigma-Aldrich, Melford Laboratories Ltd., BDH AnalaR®, and Fermentas, or unless otherwise stated.

##### *Reagents used in Cell Treatments*

Rapamycin and LY294002 were obtained from LC Laboratories. The InSolution$^{TM}$ Sinefungin, Fibroblast growth factor-2 (FGF-2), and insulin were purchased from Calbiochem. S-(5′-Adenosyl)-L-methionine chloride dihydrochloride (AdoMet) was purchased from New England BioLabs Inc. MG132, staurosporine, hydrogen peroxide, sodium arsenite, sorbitol, and sodium azide were purchased from Sigma-Aldrich. The concentration used for each reagent is stated in the text or figure as necessary.

##### *Expression Constructs and Recombinant Proteins*

The construction of pcDNA3.1/EE-p54-S6K2 expression plasmid has been described previously **[1]**. pcDNA3.1/EE-p54-S6K2 mutants (R475M, R477M, R479P, R475M/R477M (R2M), and R475M/R477M/R479M (R3M)) were made using a QuickChange site-directed mutagenesis kit (Stratagene) with pcDNA3.1/EE-p54-S6K2 as the template. Other truncated mutants (Deletion of 5 or 9 amino acids, Δ5 and Δ9, respectively) of p54-S6K2 were amplified by PCR using human S6K2 cDNA as a template. The amplicons were cloned into the BamH1/EcoR1 sites of the pcDNA3.1 plasmid (Invitrogen) in frame with the N-terminal EE-tag (MEFMPME) **[2]**.

pGEX-2T/GST-PRMT1, pGEX-2T/GST-PRMT3, and pGEX-2T/GST-PRMT6 were kindly provided by Dr. Tony Kouzarides (Wellcome Trust/Cancer Research UK Institute, University of Cambridge). The pcDNA3.1/myc-PRMT6 construct was kindly provided by Dr. Taras Valovka (Institute of Biochemistry, University of Innsbruck, Austria). All constructs were verified by restriction digestion and DNA direct sequencing. As described previously, His-S6K2 was purified from Sf9 insect cells and 80S ribosomes were purified from rat liver **[1, 3]**.

##### *Antibodies*

Polyclonal antibodies towards S6K2 and the monoclonal antibody to the EE-tag were described previously **[1]**. Anti-PRMT1 (07-404), anti-PRMT3 (07-256), and anti-asymmetric dimethylarginine (ASYM24) (07-414) antibodies were purchased from Upstate Biotechnology. Anti-PRMT6 antibody (IMG-506) was purchased from Imgenex. Anti-phospho-rpS6 (Ser240/244) (2215), anti-phospho-p70 S6K (Thr389) (p-T412 S6K) (9205), and anti-Lamin A/C (2032) were purchased from Cell Signalling Technology. Anti-β-tubulin (H-235) antibodies were purchased from Santa Cruz

Biotechnology, Inc., and anti-β-actin (AC15) antibodies were obtained from Sigma-Aldrich. Anti-Mono/DiMethyl Arginine (MMA/DMA) antibody [7E6] (ab412) was obtained from Abcam. Horseradish peroxidase (HRP)-conjugated anti-mouse (W4021) and anti-rabbit HRP (W4011) antibodies were purchased from Promega Corporation. The monoclonal antibody to the EE-tag was a gift from Dr. J Downward (CRUK). The monoclonal antibody to the Myc-tag was generated by Ivan Gout's laboratory.

***Cell Culture***

HEK293 and SCLC cell lines were obtained from the American Type Culture Collection (ATCC, USA), and maintained per the instruction of the supplier (humidified atmosphere, 10% $CO_2$, 37°C). HEK293 cells were grown in DMEM while SCLC cells were grown in RPMI 1640, supplemented with 10% heat-inactivated FBS (Hyclone), 2 mM L-glutamine, 50 U/ml penicillin, and 0.25 μg/ml streptomycin. General cell culture reagents were acquired from PAA Laboratories GmbH.

***Establishing Tetracycline-Inducible p54-S6K2 Cell Lines***

Tetracycline-inducible p54-S6K2 cell lines were generated using the Tetracycline-Regulated Expression System for mammalian cells (T-Rex System, Invitrogen) as described previously **[4]**. Briefly, the cDNA sequences of p54-S6K2 or that of mutants, with the N-terminal EE-tag, were cloned into pcDNA4/TO-inducible vector and transfected, using ExGen 500 transfection reagent (Fermentas), in T-Rex-HEK293 for 48 hours. Transfected cells were selected in complete DMEM supplemented with 5 μg/ml blasticidin (Sigma-Aldrich) and 100 μg/ml zeocin (Sigma-Aldrich). Cells were screened for tetracycline-regulated protein expression. Cells were maintained in complete DMEM supplemented with 1 μg/ml blasticidin. To induce protein expression, cells were treated with 1 μg/ml tetracycline (Sigma-Aldrich) and incubated for 24 hrs at 37°C prior to experiments.

***DNA and siRNA Transfections***

For DNA transfection studies, cells were seeded at $5x10^6$ cells per well in a 6-well plate and transfected using ExGen 500 transfection reagent (Fermentas) as recommended by the manufacturer's instructions. Cells were harvested and lysed for analysis by immunoblotting and immunoprecipitation after 24-48 hours of transfection.

***Starvation-Induced Cell Death Assay***

T-Rex-HEK293 cells stably expressing EE-S6K2 (WT), R2M mutant, or empty vector (pcDNA3.1) were plated in 12-well plates ($2 \times 10^4$ cells/well) and cell death was induced by serum starvation for 24 hrs. The percentage of dead cells was measured by the method of trypan blue exclusion **[5]**.

***Cell Lysis and Subcellular Fractionation***

Cultured cells were rinsed once with ice-cold phosphate-buffered solution (PBS) and scraped into cell lysis buffer (20 mM Tris-HCl [pH 7.5], 50mM NaF, 150 mM NaCl, 5mM EDTA [pH 8.0], 1% TritonX-100, protease inhibitors mix (Roche)). After incubation on ice for 30 min, whole-cell lysates were centrifuged (10,000*g*, 30 min, 4°C) to pellet cellular debris. Protein concentrations in the cleared lysates were measured using the Bradford assay. A cytoplasmic fraction was prepared by suspending cultured cells in ice-cold hypotonic buffer (20 mM HEPES [pH 7.9], 0.5 mM DTT) supplemented with a protease inhibitor cocktail for 15 min. This was followed by the addition of 40 µl/ml 10% Nonidet P-40 (NP-40) and 10 sec vortexing. After centrifugation (800*g*, 5 min, 4°C) to pellet the nuclei, the supernatant was subjected to centrifugation (10,000*g*, 15 min, 4°C) to collect the cytoplasmic fraction. To generate a nuclear fraction, nuclei were collected and washed once with hypotonic buffer supplemented with 240 µl/ml 10% NP-40 (800*g*, 5 min, 4°C). This was followed by a second wash in a hypotonic buffer alone. Nuclear pellets were lysed in ice-cold cell lysis buffer (20 mM Tris-HCl [pH 7.5], 150 mM NaCl, 50mM NaF, 5mM EDTA [pH 8.0], 1% Triton X-100, and a protease inhibitors cocktail). After incubation for 30 min at 4°C, the nuclear fractions were separated from the insoluble nuclear matrix material by centrifugation at 10,000*g* for 30 min at 4°C.

##### *Immunoprecipitation*

Equal amounts of total protein from the generated whole-cell lysates or fractions were incubated with the specific antibody and 50% Protein G Sepharose suspension for 3 hrs. or overnight on a rotating wheel at 4°C. Beads were washed three times in ice-cold lysis buffer (450*g*, 5 min, 4°C), prior to the elution of bound immune complexes by boiling for 5 min in 2X Laemmli sample buffer. The eluted immune complexes were subjected to SDS-PAGE and analysed by immunoblotting using specific antibodies.

##### *Immunoblotting*

SDS-PAGE-separated proteins were transferred onto 0.45-µm-pore-size PVDF membranes (Millipore) using Trans-Blot (Bio-Rad). The membranes were blocked in 5% zero-fat-dried milk in Tris-buffered saline containing 0.1% Tween 20 prior to the incubation in primary antibodies for 1-3 hrs at room temperature or overnight at 4°C. Membranes were subjected to several washes in Tris-buffered saline containing 0.1% Tween 20 (TBST) to remove excess antibodies. Immunoreactive proteins were detected by incubation with species-specific horseradish peroxidase-conjugated secondary antibody for 1 hr at room temperature. The immune complexes were detected with the Amersham Enhanced ChemiLuminescence system prior to scanning using the Fluor-S Max MultiImager system (Bio-Rad).

##### *Expression of GST-fusion Proteins*

For the expression of GST-fusion proteins (GST-PRMT1, GST-PRMT3, and GST-PRMT6) in bacteria, fusion proteins were expressed in BL21 (DE3) cells. Expression was carried out at 25°C for 3 hrs in the presence of 0.5 mM IPTG. Bacteria were centrifuged at 5000 rpm at 4°C for 20 min. prior to washing in ice-cold PBS and lysis in lysis buffer (20mM Tris-HCl [pH 7.5], 50mM NaF, 150mM NaCl, 5mM EDTA pH 8.0, 1% Triton, and protease inhibitor mixture) for 30 min on ice. Mild sonication on ice was carried out for lysates prior to centrifugation for 30 min at 18000 rpm at 4°C.

Glutathione-Sepharose 4B beads (Amersham Biosciences) were used to batch-purify GST-fusion proteins from the supernatant at 4°C for 2 hrs. Beads were washed several times in ice-cold lysis buffer at 4°C. GST-fusion proteins were either retained on beads or eluted by competition with 20 mM glutathione in 100 mM Tris-HCl [pH 8.0] and 150 mM NaCl. The eluted fusion proteins were subjected to dialysis against dialysis buffer 1 (50 mM Tris-HCl [pH 7.5], 150 mM NaCl, 1mM DTT) overnight at 4°C, and then against dialysis buffer 2 (buffer 1 plus 50 % glycerol) for 3 hrs at 4°C.

***GST Pull-Down Assay***

GST-PRMT1, GST-PRMT3, GST-PRMT6, or GST alone coupled to Glutathione-Sepharose 4B beads (Amersham Biosciences) were incubated with recombinant His-S6K2 in binding buffer (20 mM Tris-HCl [pH7.5], 150 mM NaCl, 1 % Triton, 1%BSA, 0.5% NP-40) for 3 hours on a rotating wheel at 4°C. The beads were then pelleted by centrifugation at 800*g* for 2 min and washed in binding buffer for 4 times. Complexes were boiled in 2X Laemmli sample buffer for SDS-PAGE analysis.

***In Vitro Methylation Assay***

The reaction of the *in vitro* methylation assay was carried out using recombinant GST-PRMTs and His-S6K2, the methyl donor [$^3$H]AdoMet (3 μCi) (Amersham Biosciences), and in the presence or absence of the methylation inhibitor InSolution$^{TM}$ Sinefungin (500 μM). Reactions were typically performed in 40 μl of 50 mM sodium phosphate buffer (pH 7.3) at 30°C for 3 hrs with gentle shaking. The reaction products were resolved by SDS-PAGE. Gels were stained with Coomassie brilliant blue and destained overnight in 10% (v/v) methanol and 5% (v/v) acetic acid prior to the use of EN$^3$HANCE (Amersham Biosciences) for 1 hr. To precipitate the fluor in the gel, cold water is added with gentle agitation for 30 min. Gels were dried and exposed to Fuji medical X-ray film for 48 hrs. at -70°C and labelled proteins were visualized by fluorography.

***In Vitro Kinase Assay***

HEK293 were transfected with Myc-PRMT6 and EE-S6K2 expression plasmids. Cell extracts were subjected to immunoprecipitation with either anti-EE-tag or anti-Myc-tag antibodies following treatment for 30 min with the signal transduction inhibitors; 100nM Rapamycin, 50μM LY294002, or vehicle (DMSO). The immune complexes were washed three times with lysis buffer followed by a single wash with kinase assay buffer (50 mM HEPES [pH 7.5], 10 mM $MgCl_2$, 1 mM dithiothreitol, 10 mM β-glycerophosphate). The kinase reaction was initiated in 25 μl of kinase assay buffer supplemented with 1 μM protein kinase A inhibitor (Calbiochem), 50 μM ATP, 5 μCi of [γ-$^{32}$P] ATP (Amersham Biosciences), and 20 μg of 80S ribosomes isolated from rat liver as a substrate (source of S6 protein (rpS6)) **[1, 3]**. The reaction was carried out at 30ºC for 30 min and terminated by the addition of 5X SDS-PAGE sample buffer and boiling for 10 min. Proteins were resolved by SDS-PAGE. The amount of [γ-$^{32}$P] incorporated into proteins was assessed by Fujifilm FLA-2000 phosphoimager apparatus (Bio-Rad). Control IP was achieved by incubating lysates with protein A-Sepharose beads.
